# Distinct mechanisms mediate dopamine-octopamine opponency in an insect model of olfaction

**DOI:** 10.64898/2025.12.23.696261

**Authors:** Yelyzaveta Bessonova, Ivy Clark, Ryan Sumida, Jacob Kelley, Ishaan Alva, Barani Raman

**Affiliations:** Department of Biomedical Engineering, Washington University in St. Louis; Department of Neuroscience and Behavior, University of Notre Dame

## Abstract

Neuromodulators play a key role in determining how an organism processes and responds to sensory cues. Often, different neuromodulators act on the same set of neural circuits to produce diverging behavioral outcomes. In this study, we examined how dopamine and octopamine alter odor-driven neural activity in a peripheral olfactory circuit (the antennal lobe) and thereby alter behavioral responses in an opposing manner. Our results indicate that dopamine suppressed GABAergic local neuron activity during odor stimulation. Releasing the antennal lobe network from inhibition increased the principal neural activity for all odorants and led to nonspecific increases in a behavioral appetitive response. Octopamine, on the other hand, did not alter the GABAergic inhibition in the antennal lobe but still reduced the odor-evoked principal neural activity and the behavioral responses elicited by all odorants. Thus, both dopamine and octopamine used distinct mechanisms (altering recurrent inhibition vs intrinsic excitability) to mediate opponent changes in odor-evoked neural activity and behavioral outcomes.

## Introduction

An organism’s behavioral responses to a sensory cue are not fixed and could change depending on its internal physiological state.^1^ Even when the external environment remains constant, shifts in factors such as hunger^1^, reproductive status^2^, or circadian phase^3^ can reconfigure neural circuit activity and alter how the same sensory stimuli are received and processed. In locusts, for example, the scent of fresh grass (hexanol)^4^ may trigger strong food-foraging behaviors^5^ when the animal is hungry, while evoking little or no feeding response when it is satiated^1^.

Similarly, the aggregation pheromone vinylanisole^6^ may be highly effective at promoting group formation during periods when collective behavior is advantageous and in gregarious locusts, yet less so under conditions favoring dispersal and in solitary grasshoppers. This adaptive processing and response to sensory stimuli ensures that behavior output is matched to the organism’s current needs and survival priorities.

In insects, odorants are received by olfactory receptor neurons in the antenna, which transduce the chemical cues into a neural signal that is propagated to a downstream neural circuit called the antennal lobe. The antennal lobe acts as a primary processing center and comprises two types of neurons: cholinergic projection neurons (PNs) and GABAergic local neurons (LNs), both of which receive signals from olfactory receptor neurons (ORNs).^7–9^ Odorants generate distinct patterns of activation in individual projection neurons and, at a combinatorial level, encode both stimulus identity and intensity.^10,11^ In contrast, LNs primarily contribute through inhibitory modulation of PN activities.^9^ The precise role of LNs in shaping olfactory-driven behavior, however, remains to be fully understood. It is hypothesized that various neuromodulators play a crucial role in modulating the individual PN activity in the antennal lobe.^12^ Whether these neuromodulators influence PNs directly or if the observed changes result from alterations in LN function is yet to be deciphered.

Dopamine (DA) and octopamine (OA) are key neuromodulators that are linked to various behaviors, including associative learning^13,14^, feeding^15^, flying^16^, grooming^17^, and mating^18,19^. Specifically, octopamine is linked to mediating the fight-or-flight response in locusts.^15^ Previous research reports that the injection of OA into the thorax region provokes flight-like, rhythmic firing patterns in neurons,^15^ and blockage of OA resulted in flight motion impairment.^16^ In cockroaches, injection with dopamine-containing wasp venom into the central nervous system (CNS) causes prolonged and frequent grooming.^15,17^ In honeybees, it has been shown that micro-injections of octopamine to the AL can act as a substitute for positive reward (i.e., sucrose water) during olfactory associative learning.^14^ In contrast to octopamine, dopamine has been shown to induce an opposite effect on the same set of behavioral outputs. For example, studies using aversive learning paradigms in *Manduca sexta* have demonstrated that blocking dopamine signaling in the AL can prevent learned avoidance.^13^ In *Drosophila,* dopaminergic neurons have been shown to modulate neural activity in the learning center (mushroom bodies), resulting in a variety of behavioral changes.^20^ For example, dopamine release spikes in male flies during courtship and rapidly decreases when the mating needs are satisfied.^18–20^

How do different neuromodulators alter sensory processing in a neural network in a complementary fashion and thereby adapt behavioral outputs to fit the needs of the organism? In this study, we examined this issue by assessing how odor-evoked neural responses are altered by the two key neuromodulators in the antennal lobe (dopamine and octopamine), and how these changes in neural activity produce a correlated change in a behavioral output. We show that local neurons play a crucial role in how the system alters the antennal lobe output in a neuromodulator-dependent fashion. Finally, we present a simple model of dopamine- octopamine opponency to map the neural activity alterations onto the changes in behavior. In sum, our results provide a better understanding of how opponent neuromodulators could fine-tune the same circuit to adaptively generate different behavioral responses.

## Results

### GABAergic local neurons with anti-correlated odor-evoked responses in the locust antennal lobe

The sensory input from the olfactory receptor neurons (ORNs) in the locust *(S. americana)* antenna drives responses in both cholinergic projection neurons (PNs) and GABAergic local neurons (LNs).^9,21^ While the stimulus-evoked responses in PNs have been well characterized, how LNs respond to odorants and their functional diversity is yet to be fully understood.^9,22^ We began by intracellularly characterizing local neuron responses to different odorants (20 LNs from 20 locusts; 100 LN-odor combinations; hexanol, benzaldehyde, linalool, citral, and geraniol).

Consistent with prior findings, we found that LNs did not fire fully blown sodium spikes (**Fig. 1A**).^21^ We found one category of LNs that had higher spontaneous activity, but the membrane potential became transiently hyperpolarized during odor presentation (**Fig. 1A**; LNs 1 - 3). In contrast, a second group of LNs had less spontaneous activity, but the membrane potential became depolarized during odor presentation (**Fig. 1A**; LNs 4 – 6). Unlike the PN responses that vary with odor identity, the LNs maintained their response type (hyperpolarization vs depolarization) to all odorants evaluated (**Fig. 1D**).

**Figure 1.**
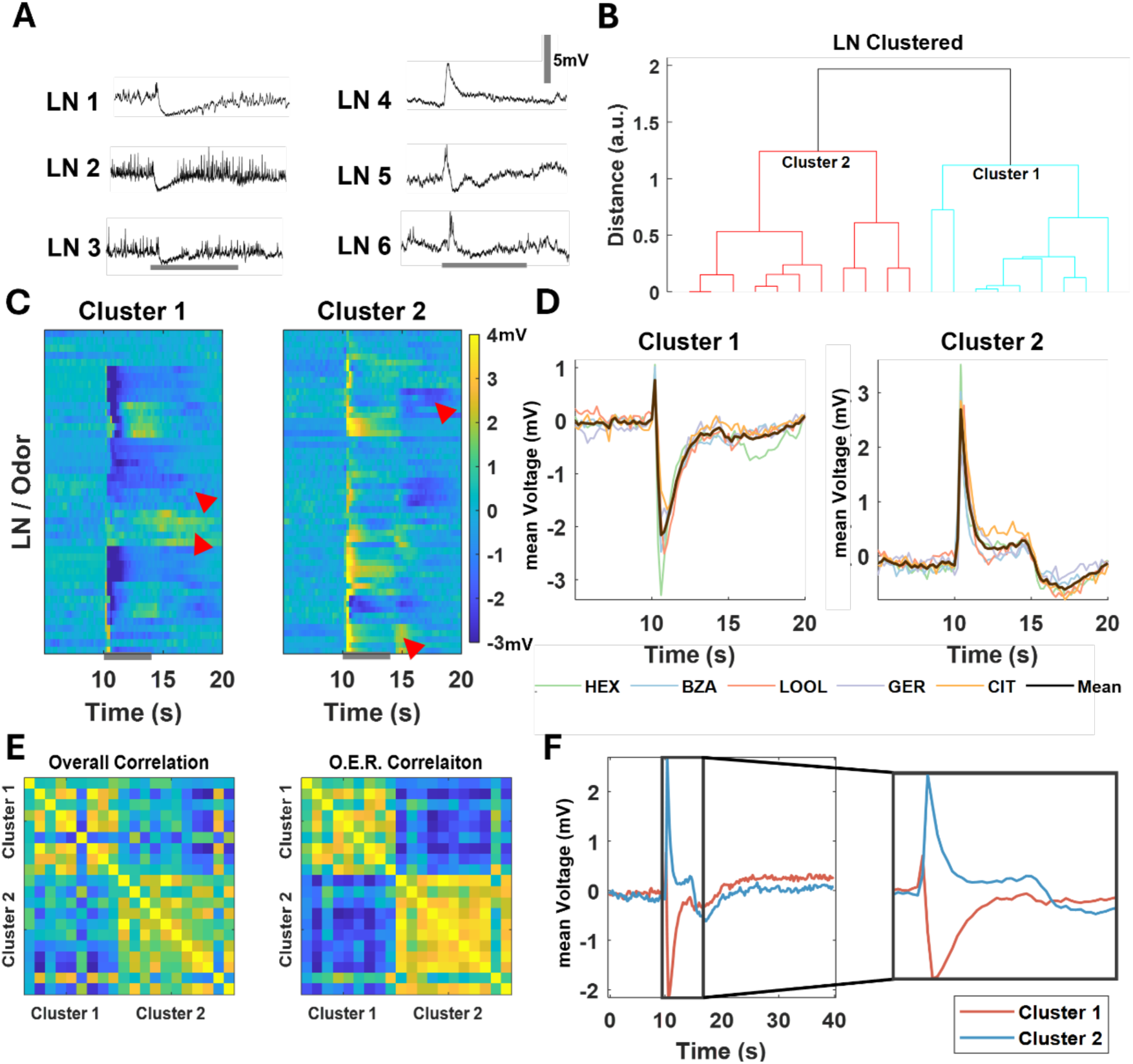
Two functional categories of odor-evoked responses in the antennal lobe local neurons (LNs). **(A)** Representative intracellular recordings show membrane potential as a function of time for six distinct local neurons (LNs) in the locust antennal lobe. Two types of odor-evoked responses were observed: high spontaneous activity and hyperpolarization during the odor presentation window (LN1-3) and low spontaneous activity and depolarization during stimulus exposure (LN4-6). The vertical grey scale bar represents 5 mV. The 4s odor presentation window is indicated by the horizontal gray bar. **(B)** The dendrogram represents the clustering of 20 local neurons into two clusters using a hierarchical clustering approach (see Methods). Cluster 1 (hyperpolarizing LNs) contains 11 cells, whereas cluster 2 (depolarizing LNs) has 9 cells. **(C)** Five-trial-averaged voltage values for each odor as a function of time are shown for both LN clusters. Responses to five odorants are shown (hexanol, benzaldehyde, linalool, citral, and geraniol; five rows for each LN). The average membrane potential in 200-ms time windows was calculated and baseline-subtracted before being shown here as a heat map. Each row corresponds to trial-averaged response to one odorant, and five consecutive rows represent one LN. The hotter color identifies depolarization, and the colder color indicates hyperpolarization. The horizontal grey bar indicates the odor presentation period. The scale bar represents the baseline-subtracted, binned, and averaged voltage in millivolts *(mV)*. **(D)** Average membrane potential across all the LNs belonging to cluster 1 and cluster 2 is shown for each odor: hexanol (*green*), benzaldehyde (*blue*), linalool (*red*), geraniol (*purple*), and citral (*yellow*). Black trace denotes the LN response averaged across all five odorants. Note that LNs in Cluster 1 have a weak and brief depolarization followed by a pronounced hyperpolarization response during odor presentation, whereas LNs in Cluster 2 showcase strong depolarization during the stimulus presentation window. These response motifs did not change much with odor identity. **(E)** Cross-correlation analysis of Cluster 1 and 2 LNs summarizing all 20 local neuron responses across all 5 odorants. Each pixel represents the correlation value between a pair of the LNs. *On the left,* the activity across all 40-second recordings was used to determine the correlation value (a measure of similarity). *On the right,* only odor-evoked response (O.E.R) was considered to calculate the correlation between pairs of LNs (both 4 seconds during and 4 seconds after odor presentation). **(F)** PSTH across all LN/odor combinations for Clusters 1 *(orange)* and 2 *(blue)* plotted together. Note *on the right,* a ten-second time window that included two seconds of pre-stimulus activity, four seconds of odor presentation, and four seconds after odor termination is shown.

We used a simple hierarchical clustering analysis (see **Methods**) to split our LN dataset into clusters or response motifs (**Fig. 1B-D**). Notably, our entire dataset of 100 LN-odor combinations split into two categories we qualitatively observed: high spontaneous activity and hyperpolarization during the stimulus period (Cluster 1: 9 LNs, 45 LN-odor combinations), and low spontaneous activity and depolarization during odor presentation (Cluster 2: 11 LNs, 55 LN- odor combinations). In both clusters, we note that few LN responses outlasted the stimulus duration, and a second bout of either excitatory or inhibitory response following odor offset was observed (**Fig. 1C**; **arrowheads**). Consistent with our interpretation that these LN responses are cell-dependent and not odor-identity-dependent, we found that mean odor-evoked activity across all LNs in each cluster was similar for all five odorants used (**Fig. 1D**). To further investigate the differences or similarities between the two clusters of LNs, we first averaged the activity across all 5 odorants for each LN. Then, we computed cross-correlation between the overall and odor-evoked activity of all LNs in both clusters (**Fig. 1E**). Cluster 1 and cluster 2 LNs were anti-correlated both before and during odor presentation.

Finally, we compared the relative timing differences in the stimulus-evoked responses following odor onset for LNs in the two clusters. We found that even the hyperpolarizing LNs had a rapid initial transient depolarization, and the peak depolarization occurred before the Cluster 2 LNs were activated (**Supplementary Fig. 1**; *right panel*). As the membrane potential voltage in Cluster 2 LNs increased, though, Cluster 1 LNs became hyperpolarized. Notably, the peak LN depolarization for many Cluster 2 LNs occurred prior to the maximum hyperpolarization epoch of Cluster 1 LNs (**Supplementary Fig. 1**; *left panel*). Finally, the hyperpolarization in the Cluster 1 LNs reduced as the activity in Cluster 2 LNs reduced (**Fig.1F**). These results suggest asymmetric, competitive interactions between the two groups of GABAergic LNs, where Cluster 2 LNs inhibit activity in Cluster 1 LNs.

Taken together, our results indicate that there are two primary cell-dependent functional categories of GABAergic LNs with anti-correlated odor-evoked responses in the locust antennal lobe.

### Dopamine alone increased the hyperpolarization of local neurons

In addition to the input from olfactory sensory neurons, the internal state of the organisms, such as hungry vs satiated, naïve vs conditioned, and/or their behavioral phenotype (i.e., solitary vs. gregarious phase), could all influence how the olfactory circuits process sensory information and therefore the overall behavioral response to an odorant.^23^ Given their broad connectivity patterns, we wondered whether the local neurons could indeed be the loci where such flexibility in olfactory processing is achieved. To study this, we next examined whether and how the local neuron responses, and therefore their function, are altered by different neuromodulators.

We began by examining how dopamine, octopamine, and serotonin perturb the spontaneous and odor-evoked responses of local neurons. We performed intracellular recordings from LNs before and after exogenous introduction of one of these neuromodulators (see **Methods**). Our results indicate that only dopamine had a pronounced effect on stimulus-evoked local neuron responses. (**Fig. 2A-B**) Enhanced hyperpolarization of the local neurons was observed in most local neurons and became amplified when the voltage traces across these neurons were averaged (**Fig. 2C-E**). Hence, these results indicate that most local neurons become suppressed or inhibited during odor presentation in the presence of dopamine.

**Figure 2:**
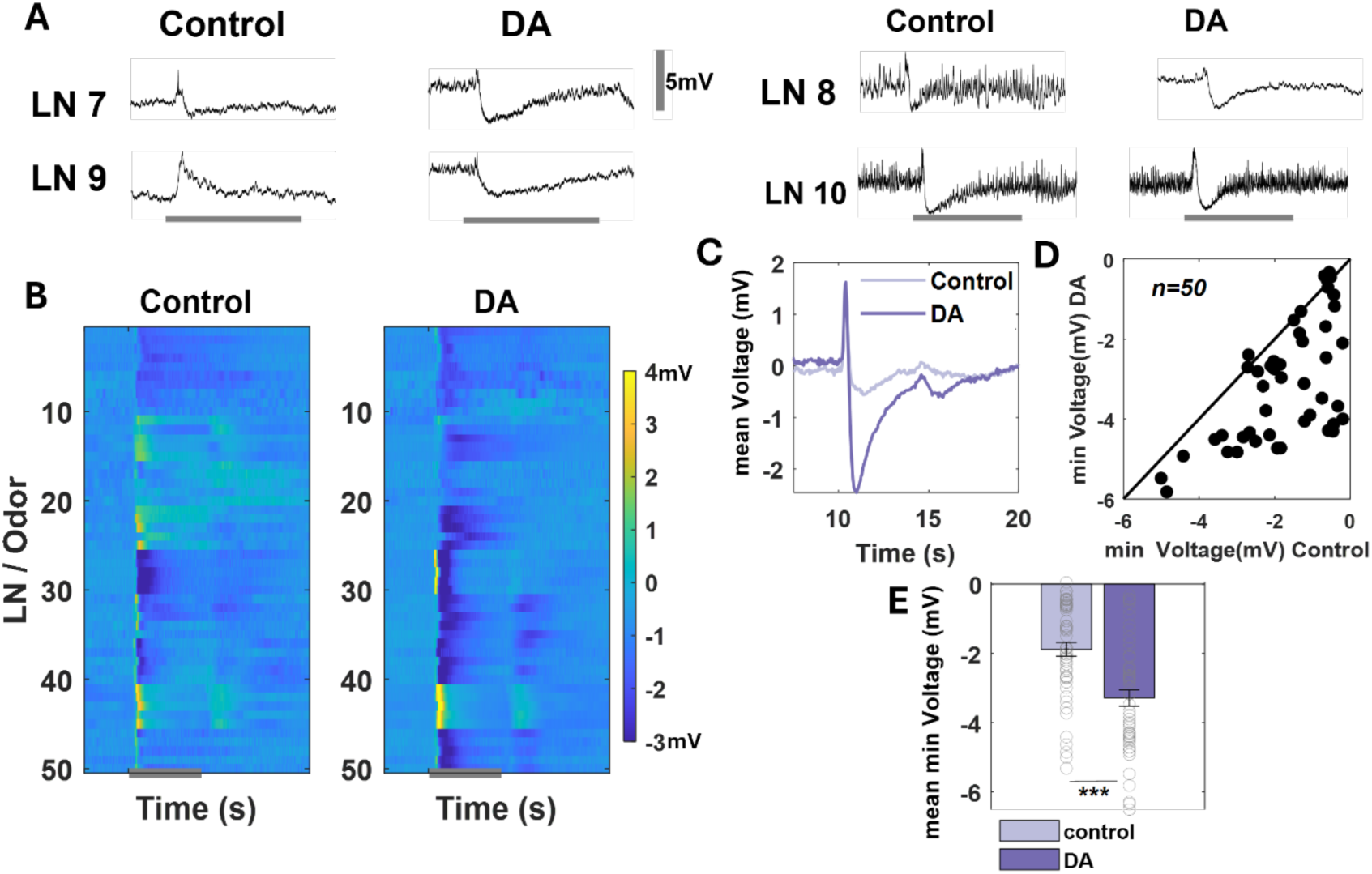
Dopamine enhances hyperpolarization in GABAergic local neurons. **(A)** Representative intracellular recording shows membrane potential as a function of time for four distinct local neurons before (control) and after application of dopamine (DA). Note that after dopamine introduction, the LN membrane potential during odor stimulation changes from depolarization to hyperpolarization. The vertical grey scale bar represents 5 mV. The horizontal grey bar identifies the odor presentation period. **(B)** Trial-averaged voltage values as a function of time are shown for the recorded LNs before (control) and after treatment with dopamine (DA). Responses to five odorants are shown (hexanol, benzaldehyde, linalool, citral, and geraniol). The raw membrane potential in 200-ms time windows was averaged and baseline-subtracted before being shown here as a heat map. Each row corresponds to trial-averaged response to one odorant, and five consecutive rows represent one LN. The hotter color identifies higher value (depolarization), and the lighter color indicates lower value (hyperpolarization). The scale bar represents the baseline-subtracted, binned averaged voltage in millivolts *(mV)*. The grey bar along the x-axis identifies the 4 seconds of odor presentation. **(C)** PSTHs of all LN/odor combinations are shown before (lighter shade of purple) and after dopamine (darker shade of purple) application. Note that DA introduction enhanced the overall hyperpolarization of the LNs. (n=50) **(D)** Minimum voltage value during odor presentation (i.e. peak hyperpolarization value) in the control condition and after neuromodulator application for each LN/odor combination. Note that all the values are below the diagonal, indicating enhancement of the hyperpolarization. (n=50) **(E)** The mean minimum voltage value across LNs during odor presentation in the control condition (light purple) and after neuromodulator application (dark purple) is shown as a bar plot. Error bars indicate SEM. Grey circles showcase minimum values for each LN/odor combination. Significant differences are identified in the plot (standard paired sample t-test,***p<0.001). (n=50)

In contrast, the majority of local neurons maintained their spontaneous and stimulus-evoked responses after application of octopamine (**Fig. 3A-B**) and serotonin (**Supplementary Fig. 4**). Interestingly, octopamine modestly increased hyperpolarization during the sustained local neuron response to a persisting odor stimulus, but reduced the local neuron hyperpolarization after stimulus termination (**Fig. 3C**, **Supplementary Fig. 4**).

**Figure 3.**
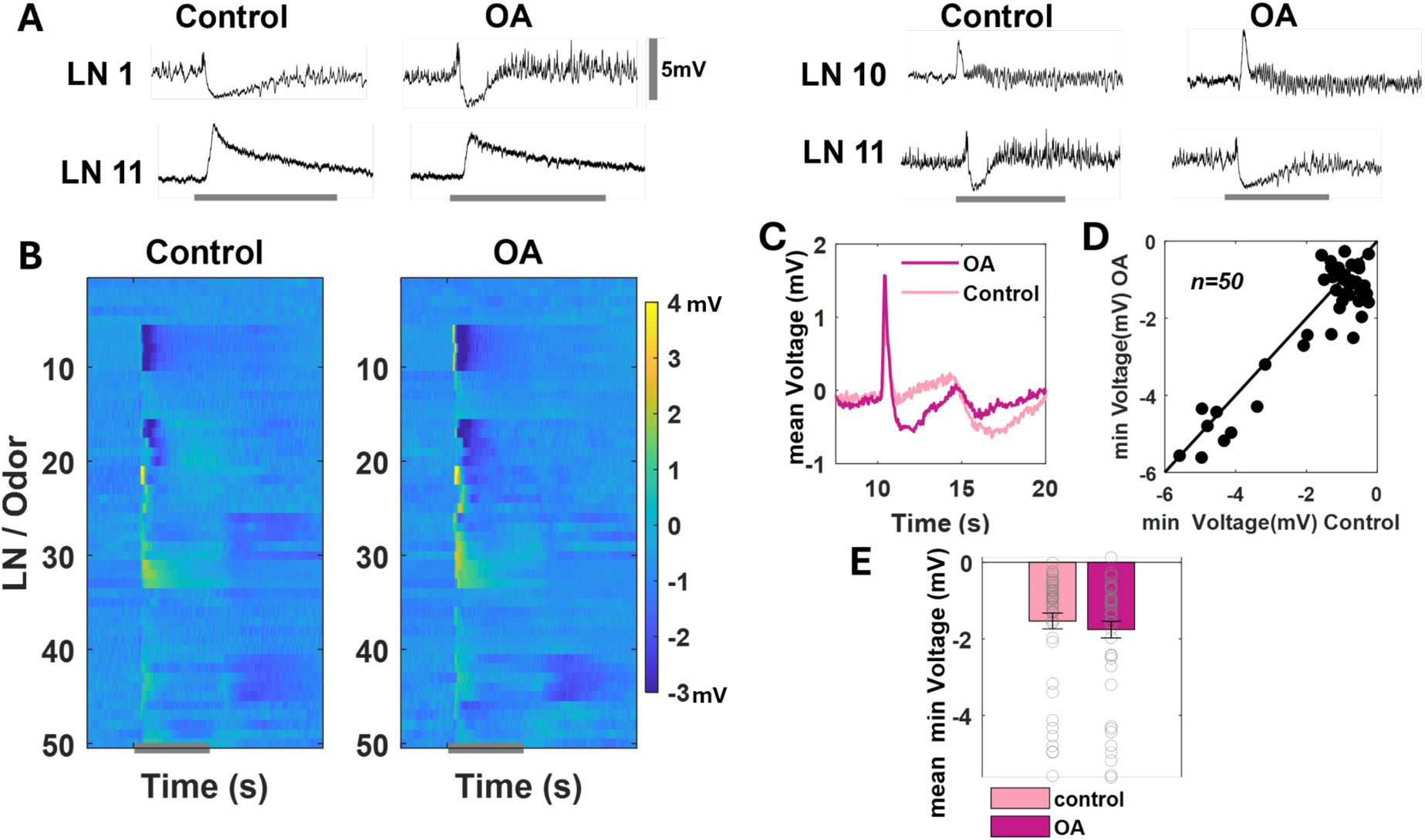
Odor-evoked responses in local neurons are maintained after octopamine introduction. **(A)** Representative intercellular recordings show membrane potential as a function of time for four distinct local neurons before (control) and after application of the octopamine (OA). No significant change in odor-evoked response was observed. The vertical grey scale bar represents 5 mV.The horizontal grey bar represents the odor presentation period. **(B)** Trial-averaged voltage values as a function of time are shown for the recorded LNs before (control) and after treatment with the octopamine. Responses to five odorants are shown (hexanol, benzaldehyde, linalool, citral, and geraniol). The raw membrane potential in 200-ms time windows was averaged and baseline-subtracted before being shown here as a heat map. Each row corresponds to the trial-averaged response to one odorant, and five consecutive rows represent one LN. The hotter color identifies higher value (depolarization), and the lighter color indicates lower value (hyperpolarization). The scale bar represents the baseline-subtracted, binned averaged voltage in millivolts *(mV)*. The grey bar identifies the 4 seconds of odor presentation. **(C)** PSTHs of all LN/odor combinations are shown after averaging. The lighter shade PSTH shows the PN responses before octopamine application, and the darker shade represents the PSTHs after OA application. (n=50) **(D)** Minimum voltage values is shown during odor presentation in the control condition and after OA application for each LN/odor combination. For OA, the values located along the diagonal indicate not much change in peak hyperpolarization after OA application. (n=50) **(E)** The minimum voltage value during odor presentation after averaging across LNs in shown for the control condition (lighter bar) and after OA application (darker bar) is shown as a bar plot. Grey circles showcase individual peak hyperpolarization values for each LN/odor combination. Error bars indicate SEM. Significant differences are identified in the plot. (n=50)

Taken together, our results indicate that amongst the neuromodulators examined, only dopamine had a strong suppressive effect on the activity of local neurons.

### Neuromodulators alter spontaneous and odor-evoked projection neuron activity in the antennal lobe

How do the observed changes in GABAergic local neuron activity modulate the cholinergic projection neurons’ output from the antennal lobe? First, we found that within seconds of dopamine application, PN activity was completely suppressed. The spiking activity gradually recovered over 2–3 minutes; however, the spontaneous activity in the network remained lower than levels observed before application of dopamine. **(Supplement Fig. 3)** This suggests that dopamine has a rapid but transient suppressive effect on PNs. No such transient effect was observed following octopamine application.

In our earlier study, we found that serotonin increased the spontaneous activity in the projection neurons and made the network activity more bursty^1^ (**Supplementary Fig. 2**). In contrast, we found here that dopamine and octopamine reduced the baseline PN activity (**Fig.4B**). For dopamine, roughly 2/3^rd^ of the PNs we recorded had lower spontaneous firings (∼20/60 cells). Octopamine suppressed the spiking activity even more and in a larger fraction of the recorded PNs (∼39/45 recorded cells). Dopamine and octopamine had opposite effects on the odor-evoked PN activity, however. While dopamine enhanced the odor-evoked responses, octopamine reduced activity in most neurons (**Fig. 4C**).

**Figure 4.**
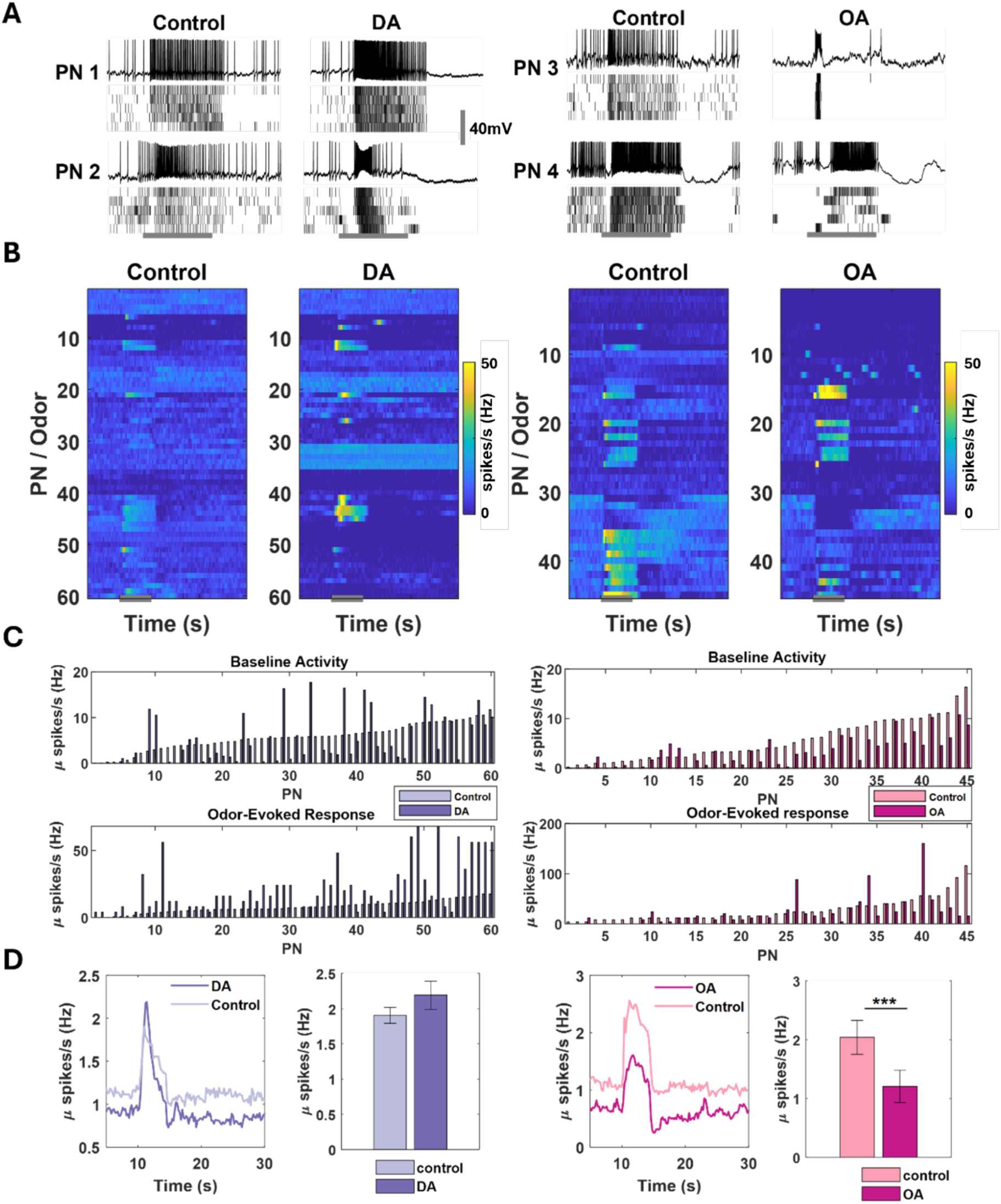
Dopamine and octopamine have opposite effects on odor-evoked projection neuron responses. **(A)** A representative intracellular recording displays the membrane potential over time for four distinct projection neurons (PNs) in the locust antennal lobe before (control) and after neuromodulator (DA – left panel, or OA – right panel) application. The raw trace for the first trial is followed by five consecutive trials presented as raster plots below the voltage trace. The horizontal gray bar indicates the 4-second odor delivery period, and the vertical gray scale bar represents 40 mV. **(B)** Trial-averaged firing activity as a function of time are shown for PNs before (control) and after treatment with dopamine (DA) or octopamine (OA). Responses to five odorants are shown for each PN (hexanol, benzaldehyde, linalool, citral, and geraniol). The firing rate in 200-ms time windows was binned and then averaged. Each row corresponds to the average of five trials and represents one PN-odor combination. The hotter color indicates a higher firing rate. The scale bar represents mean firing rate. The grey bar identifies the 4 seconds of odor presentation. **(C)** Comparison of mean firing rate during baseline or no stimulus period is shown as a bar plot for each individual PN recorded. The bars in lighter shade indicate spiking activity in the control period (i.e. no application of neuromodulator). The darker shaded bars show the mean spontaneous spiking activities in the same set of PNs after application of either DA (left panel) or OA (right panel). **(D Left)** *Left panel:* PSTH across all the PN/odor combinations before and after application of DA. The lighter shade of color represents the control, and the darker shade is the PSTH averaged across PNs after DA application. Peak odor-evoked firing rate after averaging across all PNs is shown for control (*lighter shade*) and after DA application (*darker shade*). (n=60) **(D Right)** Similar plot as in panel E1, but comparing the PSTHs and peak odor-evoked responses across PNs before and after application of OA.(n=50)(standard paired sample t-test,***p<0.001).

To examine whether these changes were odor-specific, we extracellularly recorded from a larger number of PNs and from a larger panel of odorants. We observed that the changes we observed in our intracellular recordings were also present in the extracellular dataset. The baseline or spontaneous activity in the antennal lobe network was reduced after application of dopamine or octopamine. Dopamine application increased stimulus-evoked responses for all odorants used in the panel (**Fig. 5A-B**). And octopamine reduced odor-evoked responses in the AL network.

**Figure 5.**
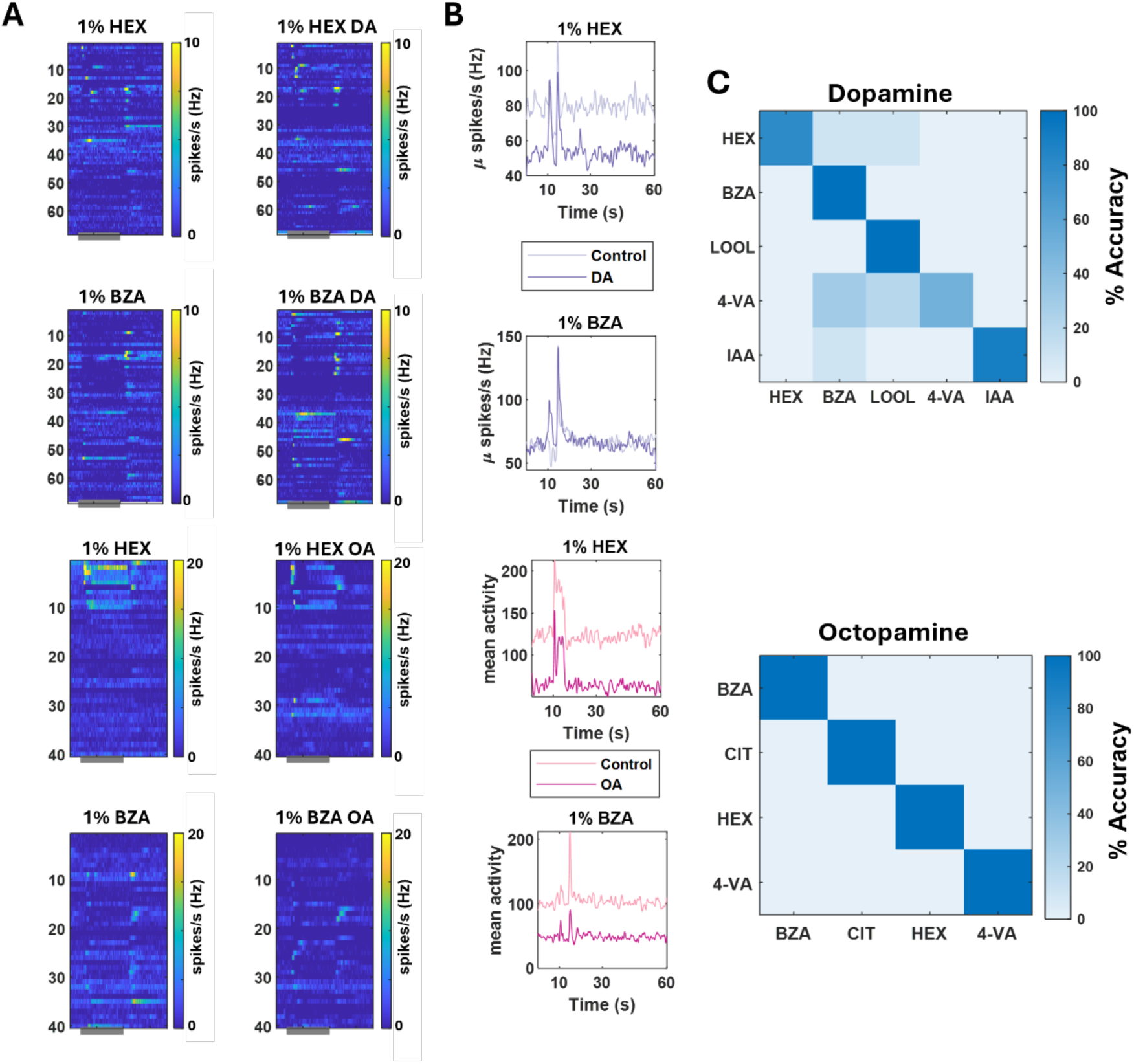
Robust odor recognition after dopamine or octopamine application. **(A)** Trial-averaged spiking activity of individual PNs (rows) is shown as a function of time for the control condition as well as after the application of DA (top panels) or OA (bottom panels). The horizontal grey bar indicates the odor presentation period. Hotter colors represent a higher firing rate**. (B) PSTHs** averaged across all of the PNs are shown as a function of time for two odorants: HEX and BZA. PSTHs in a *lighter shade* summarize PN activity before neuromodulator application, and PSTHs in a *darker shade* reveals the PN activity after DA or OA application. **(C)** Confusion matrices summarizing the classification performance of a nearest neighbor classifier is shown. The PN ensemble activity in control condition (i.e. before application of DA or OA) was used as the training response templates to be pattern matched. Each trial after DA or OA application was compared with the training templates using Cosine distance metric. The trials were assigned the same odor label as the best matching training template. Note that the confusion matrices were mostly diagonal indicating good performance and stability of ensemble neural activity in allowing robust recognition.

We wondered whether the changes in the odor-evoked PN activity altered the ensemble response fingerprint for the odorant, confounding subsequent recognition of the same odorant. To examine this issue, we did a classification analysis (see **Methods**) where the odor-evoked responses in the control case (i.e., before introduction of neuromodulators) were used as templates to recognize the stimuli (**Fig. 5C**). Our results indicate that although both dopamine and octopamine did alter responses in individual PNs the overall response fingerprint across the ensemble of PNs remained relatively consistent to allow robust recognition.

### Dopamine and octopamine have opposite effects on the palp-opening response behavior

How do changes in neural activity alter the overall behavioral outcomes? Our previous studies have shown that the projection neuron activities in the antennal lobe are a good indicator of an appetitive palp-opening response.^5^ In this behavioral assay, puffing food-related odorants such as hexanol (hex) resulted in the opening of sensory appendages close to the mouth (called ‘maxillar palps’). The palps are typically opened to touch and accept food, such as a blade of grass. Presenting non-appetitive odorants, such as benzaldehyde (bza), resulted in a weaker palp-opening response (lower palp-opening probability) compared to those observed during hexanol presentations. Therefore, we sought to understand how the changes in the antennal lobe responses we observed with the three neuromodulators alter overall palp-opening responses (PORs). In this assay, we exposed each locust to the same odorant panel as in the electrophysiological experiments (n = 20 locusts; 5 odorants: *hex, bza, lool, ger, cit, and a control: p-oil*). Each odorant was puffed ten times (**Fig. 6A**), and the probability of palp-opening response across trials and locusts was calculated for each odorant. Consistent with our prior results, hexanol had the highest probability of eliciting a POR response, and the control paraffin oil generated the weakest POR responses.

**Figure 6.**
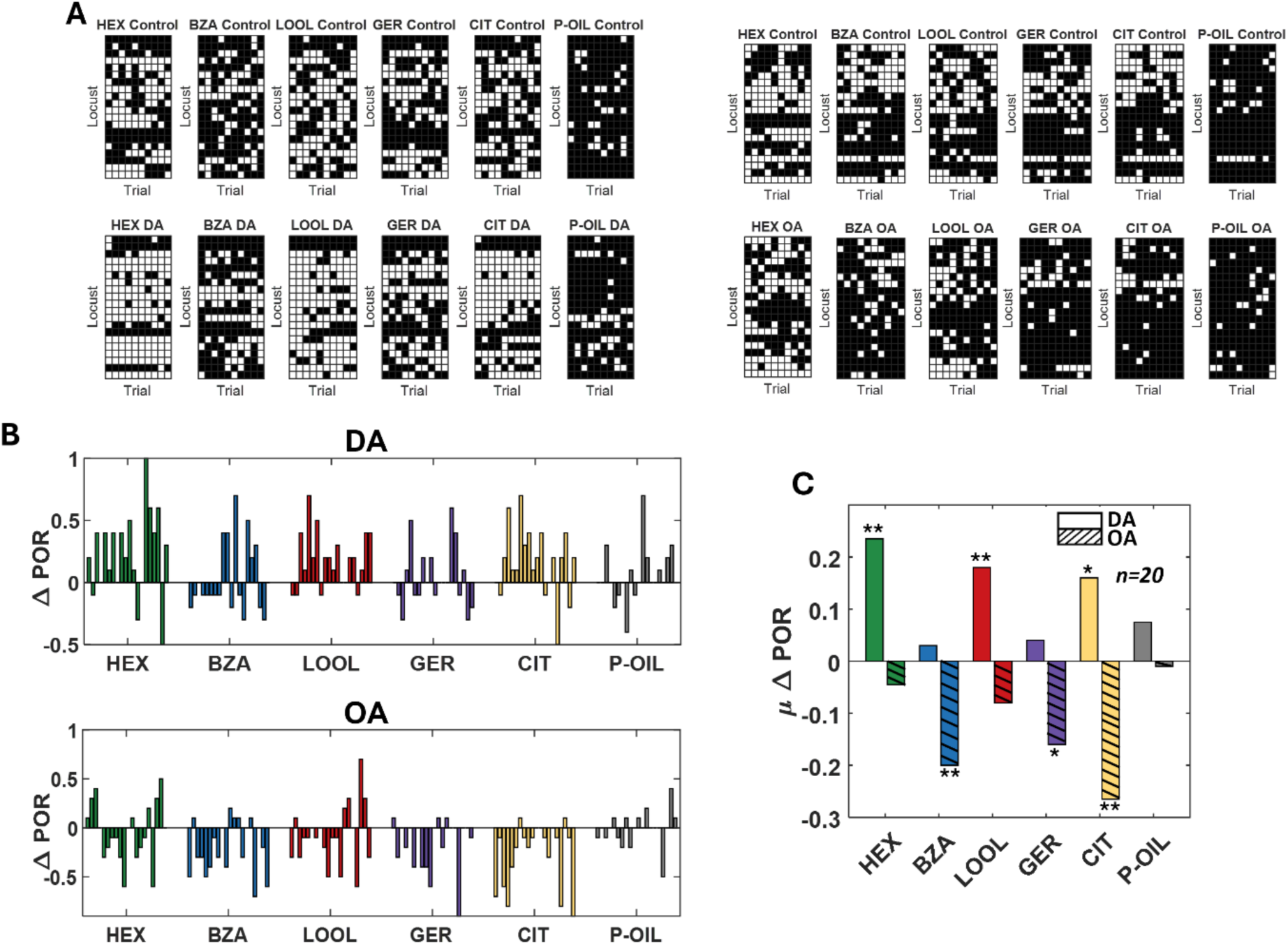
Dopamine increases and octopamine decreases the palp-opening response (POR). **(A)** A summary of trial-by-trial PORs for each locust is shown. Each trial was categorized as either POR present (white box) or absent (black box). Each row represents the PORs recorded from a single locust, and each column indicates a specific trial. PORs of twenty locusts per neuromodulator were recorded and summarized as a response matrix. The POR response matrix for the same set of locusts before and after dopamine (*on the left*) and octopamine (*on the right*) injection is shown to allow comparison. **(B)** The delta of trial-averaged PORs before and after DA (*top*) and OA (*bottom*) injections is shown for each locust as a bar plot for all four odorants in the panel. Odor panel: hexanol (green), benzaldehyde (blue), linalool (red), geraniol (purple), citral (yellow), and paraffin oil (solvent, grey). **(C)** The averaged delta values for both DA (plain) and OA (stripped) are shown for each of the odorants. Significant differences are identified in the plot (one-tailed paired-sample t-test; *p<0.05; **p<0.01; standard paired sample t-test).

We first examined the POR responses in the same set of locusts before and after dopamine injection (see **Methods**). In general, we found that the POR responses increased for all five odorants presented (**Fig. 6A**). The increase in POR was particularly significant for hexanol, linalool, and citral (**Fig. 6B, C**). A small but insignificant increase in POR was observed for the other two odorants. Next, we used the same POR assay and the odor panel on a different set of locusts to examine the effects of octopamine on the overall appetitive behavioral responses (**Fig. 6B, C**). In contrast to dopamine, we found that injection of octopamine reduced the overall POR responses to all five odorants examined. This reduction was significant in benzaldehyde, geraniol, and citral. The non-odor-specific increase and decrease we observed with octopamine and dopamine are in contrast with our earlier results that showed serotonin increased and decreased POR responses in an odor-specific manner.^1^ Notably, the opposite effect observed for dopamine and octopamine correlated well with changes in the odor-evoked neural activity we observed in the antennal lobe.

### A simple circuit model for achieving DA and OA opponency

Our results indicated that the application of dopamine increased odor-evoked PN responses and behavioral palp-opening responses to all odorants. In contrast, octopamine introduction decreased odor-evoked responses and PORs. Furthermore, dopamine alone increased the hyperpolarization and suppression of GABAergic local neuron responses in the antennal lobe circuit. We wondered whether there was a simple model that could account for all our observations.

In our recent study to understand how serotonin alters neural activity and behavior, we found that serotonin increased odor-evoked responses elicited by all odorants, but PORs increased or decreased in an odor-specific manner.^1^ A statistical model that predicted behavioral POR from neural activity indicated the need for two groups of PNs, one to increase PORs and another to suppress the same behavioral response.

Our results here indicate two different types of LNs: one group with lower spontaneous activity that depolarized during odor presentations, and the second group that had higher spontaneous activity but became hyperpolarized during odor presentations. A simple antennal lobe model integrating competitive PN and LN networks is shown in **Fig. 7A**. Increased activity in PN ensemble 2 increases POR behavior, whereas activation of PN ensemble 1 suppresses the same behavioral outcome. The two categories of LNs mediate competitive interactions between these two groups of PNs.

**Figure 7.**
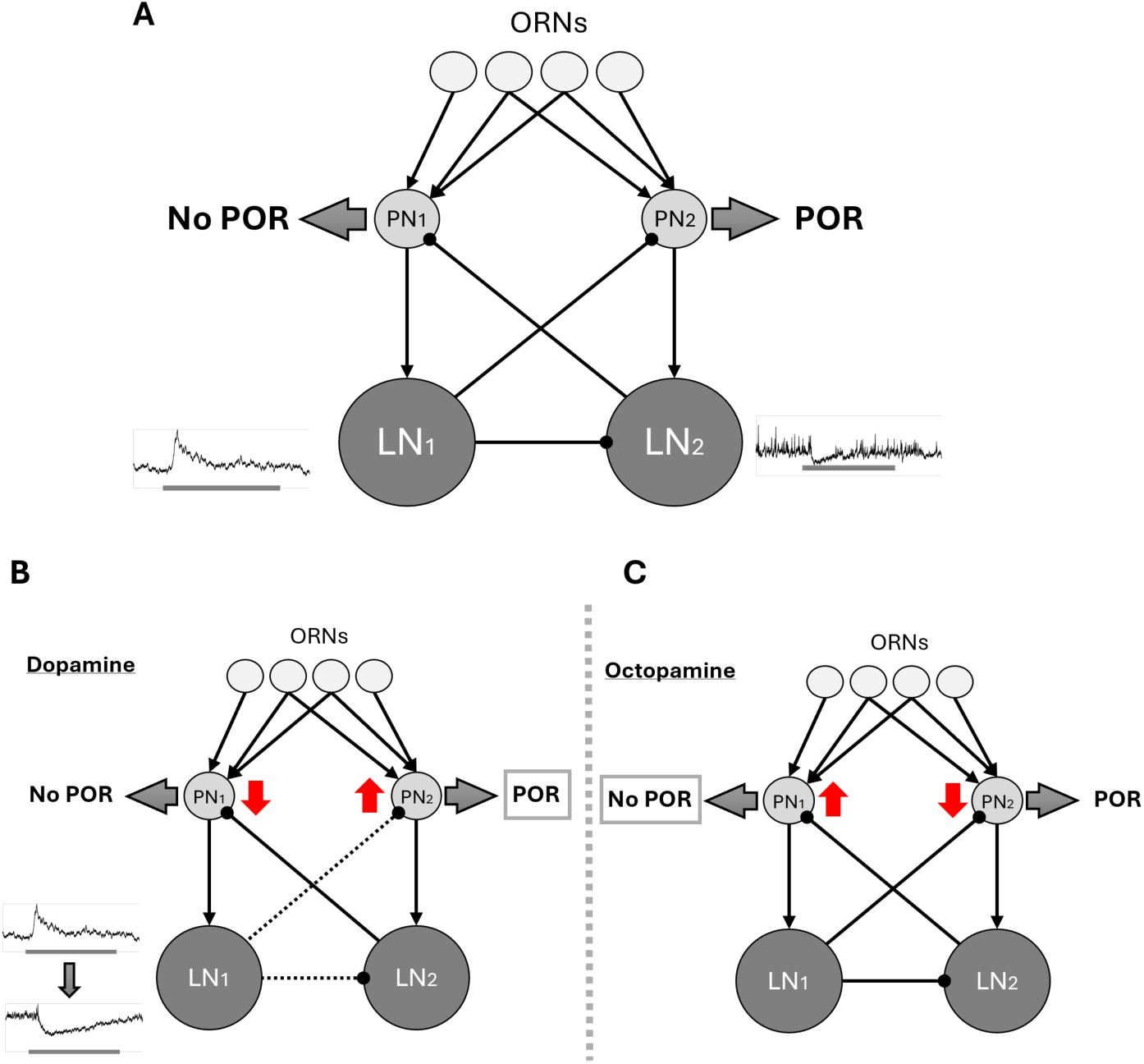
A simple competitive neural network for modulating neural-behavioral mapping. **(A)** A simplified schematic of the olfactory circuit components in the locust antennal lobe. In the figure, three types of neurons are presented: ORNs (white), PNs (light grey), and LNs (dark grey). PNs and LNs are divided into 2 groups according to the type (1 or 2). Each connection between the neurons represents excitation (arrow) or inhibition (ball arrow). In this model, type 1 PNs are responsible for suppressing PORs (no POR group), whereas type 2 PNs are responsible for increasing PORs (POR group). In detail, ORNs provide excitation to PNs and LNs (note that only ORN-PN connection shown in the schematic), each category of PNs provide excitation to one group of LNs, and LNs provide inhibition to opposite PN groups and thereby mediate competitive interactions in the network. Note, LNs type 1 are the depolarizing type of LNs that also provide inhibition to LNs type 2. The overall behavioral outcomes are determined by the overall neural responses in these two categories of PNs. **(B)** A schematic of the effect of dopamine on the antennal lobe neural circuit. The dashed line represents a decrease in the inhibition from LN group 1 to the other LN group and to the PN ensemble 2. The release in inhibition will result in increased PN ensemble 2 activity and therefore result in increased behavioral PORs for all odorants. The upward red arrow represents an increase in neural activity, whereas the red downward arrow means a decrease. **(C)** Octopamine does not alter LN activity. Therefore, the only other way to decrease the odor-evoked PN responses in all PNs, as experimentally observed, is by reducing the ORN input to both categories of PNs or directly altering PN excitability.

As we noted earlier, dopamine hyperpolarized and suppressed responses of the depolarizing LNs. In the model presented, suppression of this LN category released PN ensemble 2, which drives POR behavior and therefore would increase the overall PORs to all odorants, as we observed in our behavioral experiments. (**Fig. 7B**) Octopamine, on the other hand, did not significantly change the local neuron activity. However, the overall odor-evoked responses decreased in most PNs we recorded. Therefore, either the olfactory receptor neuron input that drives responses in PNs or PN excitability directly must be altered by this neuromodulator. Such a reduction in input drive or excitability will reduce the odor-evoked responses for both categories of PNs and reduce the overall behavioral responses to all odorants. (**Fig. 7C**)

## Discussion

In this study, we investigated how different neuromodulators could act on a single olfactory neural circuit and alter behavioral outcomes in opposing ways. Our results indicate that dopamine hyperpolarized GABAergic local neurons and disinhibited cholinergic projection neurons in the antennal lobe. As a result, the overall projection neuron activity, which can be regarded as an indicator of the overall output of the antennal lobe activity, increased in a non-odor-specific fashion. In contrast, octopamine did not significantly alter the inhibitory neuron activity. Nevertheless, our data indicate that both the spontaneous and odor-evoked projection neuron responses were reduced after octopamine application. The reduction in the odor- evoked projection neural activity correlated with the non-specific reduction in an appetitive behavioral response. In sum, dopamine and octopamine brought about opposite behavioral effects, but these changes were mediated by distinct neural mechanisms.

Dopamine and octopamine have been implicated in several important insect behaviors, including aversive and appetitive learning^24,25^, short-term and long-term memory formation^26,27^, feeding^28,29^, and flying and grooming^30^. Often, DA and OA have opposing effects on the same behavior. In honey bees, dopamine has been shown to substitute as negative reinforcement in place of electric shock during aversive conditioning^31,32^. Whereas, the release of octopamine in the honey bee antennal lobe or mushroom body calyces is known to be a suitable replacement for sugar water reward during appetitive conditioning^31,33^. Our results reveal the opponent action of dopamine-octopamine in modulating odor-evoked innate appetitive responses. While octopamine reduced innate odor-evoked PORs, dopamine application increased the PORs for all odorants. These non-stimulus-specific changes in behavioral outputs following octopamine and dopamine introductions are in stark contrast to our earlier serotonin results^34^. Exogenous introduction of serotonin increased or decreased the palp-opening responses in an odor- identity dependent manner. Together, these results indicate that these three key neuromodulators can alter the odor-behavior mapping in a non-congruent manner.

What are the neural mechanisms underpinning this octopamine-dopamine opponency? Earlier results in both vertebrates^35–37^, invertebrate^38–41^, and nematode^42^ models of olfaction have shown that olfactory sensory neuron input to the olfactory bulb or antennal lobe could be altered by dopamine (vertebrates) and octopamine (invertebrates). In vertebrates, dopamine is known to reduce the sensory neuron input onto the olfactory bulb neurons^35,37^. Further, dopamine receptors are co-expressed with GABAergic receptors in inhibitory neurons that mediate feedforward and recurrent inhibitory interactions within the olfactory bulb^35,43^. In invertebrates, while dopaminergic modulation of sensory neuron activity is not well understood, in vitro studies have revealed that dopamine indeed modulates the excitability of antennal lobe neurons^44^. Further, introducing dopamine in the moth antennal lobe increased the odor-evoked responses for all odorants^45^. Our results are consistent with these earlier studies and reveal that dopamine not only gates the overall inhibition level in the antennal circuits, but also that dopamine-mediated disinhibition of antennal lobe circuits increases odor- evoked projection neurons’ activity and appetitive behavioral response to all odorants.

Octopamine, on the other hand, is known to selectively enhance the olfactory receptor neuron responses to insect pheromone odorants^40,41^. The changes in olfactory sensory neuron activity have been suggested to involve alterations in the secondary messenger levels in olfactory receptor neurons^46^. In the downstream antennal lobe, octopamine introduction had a mixed effect, with both increases and decreases in odor-evoked responses being observed^47^. Our results differ from these earlier studies and show that octopamine did not alter the odor-evoked responses of GABAergic local neurons but significantly reduced the overall spontaneous and odor-evoked spiking levels in cholinergic projection neurons. Whether this reduction occurs due to modulation of sensory input from olfactory receptor neurons or direct alterations of the excitability of projection neurons needs further investigation. Preliminary examination of projection neuron spike shape before and after introduction of octopamine reveals that PN spike width increases after octopamine introduction (i.e., slower repolarization), and the subsequent hyperpolarization is diminished (**Supplementary Fig. 5**). The precise manner in which octopamine alters PN excitability remains to be determined.

Finally, we combined changes in physiology and behavior we observed following dopamine and octopamine introductions with our earlier results^1^ showing serotonin can increase or decrease PORs in an odor-specific manner. The simplest antennal lobe network model that integrated all our results required two groups of projection neurons: one subgroup to increase the appetitive behavioral response and a second group to reduce or suppress the same behavioral output.

Notably, two subgroups of GABAergic neurons mediated competitive interactions between the projection neuron ensembles. While dopamine influenced the level of inhibition in one group of local neurons, serotonin and octopamine must have putatively influenced the input drive or excitability of the projection neurons. This later model prediction, while consistent with prior physiological findings^39–41,43,44^, needs further careful examination.

## Author Contributions

B.R. and Y.B. conceived the study and designed the experiments. Y.B. and R.S. performed the intracellular recordings. I.C. obtained the behavioral dataset. J.K. and I.A. collected the extracellular projection neuron datasets. All authors analyzed various aspects of the collected datasets. Y.B. and B.R. wrote the paper, incorporating inputs and comments from all authors. B.R. provided overall supervision.

## Acknowledgements

We thank members of the Raman Lab (Washington University in St. Louis) and members of the Behavioral Plasticity Research Institute for their feedback on the manuscript and earlier presentation. We thank Pearl Olsen for insect care. This research was supported by NSF grant# 2021795 and 2319060, AFOSR grant# FA95502310461, and ONR grant# N00014-21-1-2343 to BR.

## Code availability

The custom code and datasets used to generate figures in this article are publicly available on GitHub: https://github.com/b0r2023/Distinct-mechanisms-mediate-dopamine-octopamine-opponency-in-an-insect-model-of-olfaction

## Declaration of Interests

The authors declare that they have no competing interests.

## Methods

### Animals

We used adult *Schistocerca americana* of both sexes from a crowded colony for our electrophysiology and behavioral experiments. Sixth-instar locusts were identified by fully developed wings and soft cuticle in the neck area.

### Odor Stimulation

The odor stimulus was delivered using a standard procedure.^10,11,21,48,49^ Briefly, all odorants were diluted to 1% concentration by volume (v/v) in mineral oil. 20 ml of diluted odor solution was kept in 60 ml sealed bottles. During stimulus presentations, a constant volume (0.1 L per minute) from the odor bottle headspace was displaced and injected into the carrier stream (0.9 L per minute) using a pneumatic pico pump (WPI Inc., PV-820). A vacuum funnel placed right behind the locust antenna ensured the removal of odor vapors.

### Odor Panel

The odorants were selected based on chemical structure and ecological relevance. Therefore, we chose a very diverse odor panel that consisted of a food odorant– hexanol^4^– and a putative aggregation pheromone – benzaldehyde^50^. In addition, we used three aversive odorants – linalool^51^ and citral^52^, and geraniol^53^– that are known to promote walking behavior in locusts.^54^ These odors were specifically selected based on our previous neural and behavioral experiments in this model.

### Behavior Experiments

Sixth instar locusts of either sex were starved for 24-48 hours before the experiment. Locusts were immobilized by placing them in a plastic tube and securing their body with black electric tape.^34^ The head of the locusts, along with the antenna and maxillary palps, protruded out of this immobilization tube. Note that the maxillary palps are sensory organs close to the mouth parts used to grab food and facilitate the feeding process. Locusts were given 20 – 30 minutes to acclimatize after placement in the immobilization tube.

Each locust was presented with one concentration of five odorants and one control (hexanol, benzaldehyde, linalool, geraniol, citral, and paraffin oil) in a pseudorandomized order. The odor pulse was 4 seconds in duration, and the inter-pulse interval was set to 60 seconds. The experiments were recorded using a web camera (Microsoft). The camera was fully automated with a custom MATLAB (MathWorks, Natick, MA) script to start recording 2 seconds before the odor pulse and end recording at odor termination. An LED was used to track the stimulus onset/offset. The POR responses were scored offline. Responses to each odorant were scored as 0 or 1 depending on whether the palps remained closed or opened. A positive POR was defined as a movement of the maxillary palps during the odor presentation window.

### Neuromodulator Treatment

After the initial set of POR experiments, a 0.1 M dopamine, octopamine, or serotonin solution was injected directly into the locust’s head. The needle of a U-100 syringe was inserted slightly under the cuticle of the locust head, ∼ 1mm above the median ocellus. 1μL of the solution was injected, and the opening was sealed with a small amount of melted dental wax. The locust was left to stabilize for 30 minutes after the injection and before the second set of behavioral experiments.

We did electrophysiology experiments 5-10 minutes after the bath application of DA or OA. A longer delay (30 min) was required for our behavioral experiments, as the locusts tended to be a bit more agitated after the injection of the solution.

### Electrophysiological Experiments Surgery

Sixth instar locusts of either sex were used for these experiments. The legs and wings were removed, and the locust was immobilized on a platform. The head was fixed with wax, and a cup was built around the head to hold a saline solution. The locust antennae were held in place using clear tubing and allowed to pass through the wax cup. The cuticle between the antennae was removed, and the air sacs/trachea were removed to expose the brain. The gut was then removed, and a metal wire platform was placed underneath the brain to lift and stabilize it.

Finally, the transparent sheath covering the brain was carefully removed using sharp forceps. Locust’s brains were super-fused with an artificial saline buffer. A visual demonstration of this entire protocol is available online.^55^

### In-vivo electrophysiology (single cell)

Intracellular recordings were performed using glass electrodes (resistance 8-15 MΩ) filled with intracellular saline (130mM L-aspartic acid potassium salt, 2mM MgCl2, 1mM CaCl2, 10mM HEPES, 10mM EGTA, 2mM Na2ATP, 3mM D-glucose, 0.1mM cAMP; osmolarity ∼315mmol/kg; pH 7.0). Glass electrodes were pulled using a micropipette puller (Sutter Instrument, Novato, CA). Spontaneous activity, as well as real-time odor-evoked responses, were recorded in the current clamp configuration for both PNs and LNs. Each set of experiments consisted of 5 consecutive 40-second trials, with a 20-second inter-trial interval. Odor stimulation was performed at the 10^th^ second of the trial for 4 seconds. Voltage signals were amplified (Axoclamp-2B, Molecular Devices) and saved using a custom MATLAB script.

After monitoring the responses to the odor panel, a neuromodulator solution was applied directly into the bath using a thin nozzle pipette. The dopamine/serotonin/octopamine solution (0.01M dopamine/octopamine/serotonin hydrochloride in locust saline buffer) was made fresh before every experiment due to its light sensitivity. The same set of recordings and odor panel was repeated 5-10 minutes following dopamine/octopamine/serotonin application. The protocol used to record from projection neurons and local neurons was identical.

### In-vivo electrophysiology (multi-cell)

Locust brains were superfused with a saline buffer and continuously perfused at ∼40 mL per hour. A silver-chloride reference electrode was then inserted into the saline. Multi-unit recordings from projection neurons (PNs) in the antennal lobe were made using a 4 x 4 silicon probe (NeuroNexus), where each contact was plated with gold prior to the start of the experiment to an impedance of 300 kiloohms. Data was acquired using one of two experimental setups. Each acquired data at 15 kHz and filtered the data between 0.3 and 6 kHz. One used a custom-built amplifier system (Caltech) providing a 10,000 gain and a custom MATLAB (MathWorks, Natick, MA) script to control experimental parameters. The other used an INTAN 32-channel amplifier and a custom Python script to control parameters. After baseline responses to the odor panel were collected, the saline in the dripline was replaced with a 0.1 mM solution of the desired neuromodulator dissolved in the same saline buffer used previously. After waiting 10-15 minutes for the neuromodulator to pass through the dripline and settle in the wax cup enclosing the brain, the second set of responses to the odor panel was collected.

### Spike Sorting

Ofline spike sorting was performed using the open-source Python library, SpikeInterface. The data was band-pass filtered between 0.3 and 3.5 kHz and whitened to eliminate noise and prepare the data for input to the spike sorter. The data was sorted using MountainSort5. To identify single units, a combination of SpikeInterface and custom quality metrics was used.

Single units had SD-Ratio < 2 noise s.d, unit cluster separation >5 noise s.d., and the number of spikes within 20 ms <15%. A final manual curation step removed any remaining noise units or unstable units.

## Analysis

### Probability of POR calculation

POR responses were scored in a binary fashion. POR responses of locusts were summed across ten trials and across all locusts. The probability for each odorant was calculated as follows:

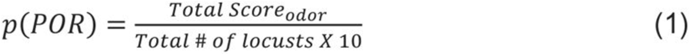

Significant differences between the POR responses observed before and after neuromodulator application were calculated using a single-tailed paired-sample *t-*test (‘*ttest’* built-in function in MATLAB).

### Local Neurons Electrophysiology Data

Altogether, we collected data from 30 local neurons with the responses to 5 odorants (Hex, Bza, Lool, Ger, and Cit), resulting in 150 LN-odor combinations. After the control recording, a particular neuromodulator was applied into the bath: dopamine, octopamine, and serotonin (10 LNs each). The electrophysiological activity was recorded intracellularly, and MATLAB (MathWorks, Natick, MA) custom code was used for analysis. Each recording was preprocessed and converted into a numerical matrix of voltage values. Due to variability in the baseline values and response dynamics, the data were normalized using MATLAB’s built-in function “normalize”. In the antennal lobe, the local neurons were identified due to their morphology (large compared to projection neurons) and lack of full-blown action potentials.

### Local Neuron Response Classification

For each LN, we compared the mean baseline membrane voltage during the pre-stimulus period (mean + 2 *standard deviation) with membrane voltage values observed during the 1-second window during odor presentation. Because the neural response was rapid and did not persist throughout the entire odor presentation, only this shorter 1-second window following odor onset was analyzed. If the mean membrane voltage value during odor presentation exceeded the pre-stimulus baseline, the LN response was classified as depolarization; if it was lower, the LN response was classified as hyperpolarization.

### Projection Neurons Electrophysiology Data

In total, electrophysiological data from 115 odor-neuron combinations were obtained intracellularly and used for analyses. Each recording was preprocessed and converted into a response matrix. The MATLAB built-in function “findpeaks” was used for identifying action potentials. In total, our dataset includes recordings from 21 PNs. All PNs were tested on a panel of five odorants (Hex, Bza, Lool, Ger, and Cit). Note that in the locust antennal lobe, only PNs fire full-blown sodium spikes. GABAergic local neurons only fire calcium spikelets and can be easily distinguished from PNs.

### Hierarchical Clustering Analysis

The same clustering method was used for both PNs and LNs. Prior to clustering, the data were separated into two matrices: before and after application of the neuromodulator. To identify patterns in the dataset, we performed hierarchical clustering using MATLAB (MathWorks, Natick, MA). Pairwise distances between neurons were computed using the correlation distance metric, which captures similarity in the shape of the data profiles rather than their absolute magnitudes. These distances were then used to construct a hierarchical cluster tree using the ’complete’ linkage method, which defines the distance between clusters as the maximum distance between their individual elements. The resulting dendrogram was used to determine cluster assignments with the ‘cluster’ function, which partitions the data into a specified number of clusters by cutting the hierarchical tree at the appropriate level. The number of clusters was selected based on visual inspection of a dendrogram. Please note that clustering was based on the hexanol results and later applied to other odor responses.

### Correlation analysis (projection neurons)

The correlation analysis was done neuron-by-neuron. Each pixel or matrix element in the neuron-neuron correlation plot indicates the correlation value between neural activity vectors during odor presentation of the *i*^th^ and *j*^th^ neuron. All neuron-by-neuron correlation analyses were computed using high-dimensional response vectors. Correlations were calculated as:

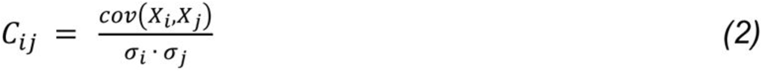

Here, *i* and *j* represent a neuron (only during odor presentation), *X_i_, X_j_* represent the neural activity vector of the *i^th^* and *j^th^* neuron, respectively, while σ*_i_*, σ*_j_* are the standard deviations of spiking activities of the *i^th^* and *j^th^* neurons.

### Classification method: (multi-unit)

Extracellular datasets comprising neural responses from sixty-eight PNs to examine the effect of dopamine and forty PNs to examine the effects of octopamine were utilized for analysis shown in (**Fig.5**). Spike counts of individual projection neurons were binned over 50 ms time window and averaged over the entire odor presentation window to get one high-dimensional PN response vector per odor per trial (dopamine dataset: 68 PNs x 10 trials per odor; octopamine dataset: 40 PNs x 10 trials per odor). Only a subset of odorants that had evoked strong and distinct responses in the control dataset (i.e., before neuromodulator application) was used in the classification analysis. This included the following odorants for the dopamine dataset: hexanol, benzaldehyde, linalool, 4-vinyl anisole, and isoamyl acetate. Similarly, only the following four odorants were utilized for classification analysis in the octopamine dataset: hexanol, benzaldehyde, 4-vinyl anisole, and citral.

A simple nearest neighbor classification approach was utilized. The PN responses before application of dopamine or octopamine (i.e., control trials) were used as training response templates to be pattern-matched (10 templates for each odorant, corresponding to 10 trials).

The PN responses after application of dopamine or octopamine were used as the testing data that needed to be classified. A cosine distance metric was used to compute the similarity of a test trial with each of the training trial ensemble PN response templates, and the odor label of the training/control trial that best matched was assigned to the test trial. The classification percentages were computed and displayed as confusion matrices.

### Action Potential Waveform Normalization and Averaging

Intracellular recordings from 20 projection neurons (12 from dopamine experiments and 8 from octopamine experiments) were used to analyze action potential (AP) waveform shape across conditions. Individual action potentials were detected using MATLAB’s “findpeaks” function.

Control data were pooled from both dopamine and octopamine control recordings.

For each detected AP, a local baseline was calculated to vertically align waveforms prior to averaging. A dual time window approach was implemented for baseline calculation: if another AP occurred within 10 ms before the current peak, the baseline was defined as the mean voltage during a short window (5-2 ms before peak); otherwise, a long window (10-2 ms before peak) was used. Each AP waveform was extracted as a 40 ms segment centered on the peak (10 ms before to 30 ms after). Waveforms were baseline-corrected by subtracting the local baseline, then normalized to unit amplitude by dividing by the peak value. The APs were normalized to avoid confounding amplitude changes due to patch clamp seal deterioration with neuromodulator-induced effects. AP waveforms were transformed as follows:

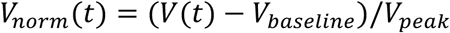

𝑉(𝑡) - raw voltage trace; 𝑉_𝑏𝑎𝑠𝑒𝑙𝑖𝑛𝑒_ - local baseline voltage𝑉_𝑝𝑒𝑎𝑘_ - peak amplitude.

Action potentials were analyzed separately during spontaneous activity (0-10 s within each trial) and odor-evoked activity (10-14 s within each trial) windows. Mean waveforms for each condition were calculated by averaging all normalized waveforms.

### Action Potential Feature Calculation

Two quantitative features were extracted from each normalized action potential waveform:

1. Full-Width at Half-Maximum (FWHM) was defined as the temporal width of the AP at 50% of its normalized peak amplitude. The half-maximum voltage was calculated as:

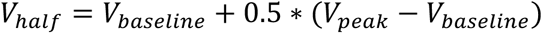

FWHM was computed as the time interval between the rising phase and falling phase crossings of this threshold.

2. Afterhyperpolarization (AHP) Depth was defined as the minimum voltage within 30 ms following the AP peak, expressed as a fraction of the peak amplitude due to normalization. To prevent contamination from subsequent spikes, the AHP measurement window was truncated at the onset of the next spike’s rising phase when another AP occurred within 30 ms.

Population-level distributions of FWHM and AHP Depth were visualized as normalized probability histograms with 30 bins.

## Supplementary Figures

**Supplementary Figure 1.**
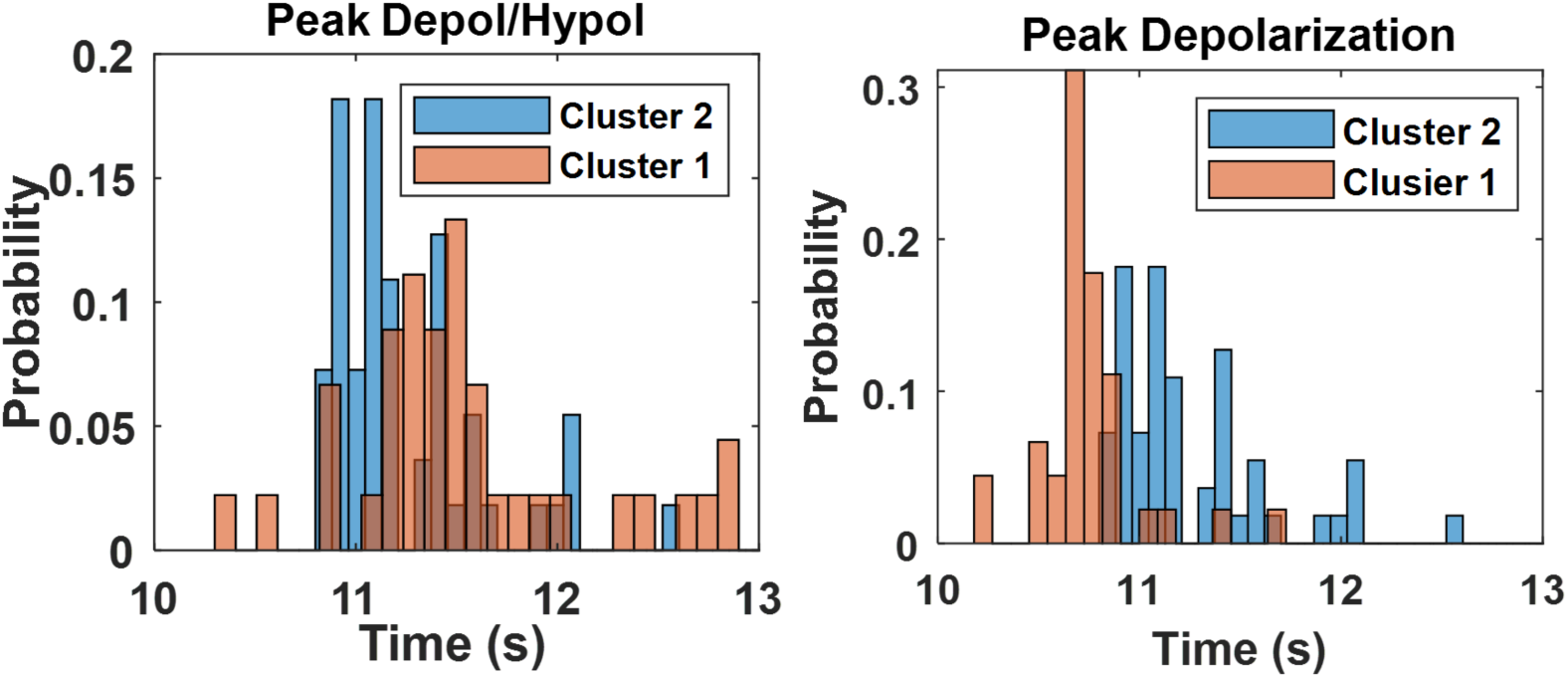
Timing differences in the peak depolarization and the peak hyperpolarization for the two categories of LNs. *On the left*, a histogram representing timing of the peak depolarization for Cluster 2 (blue) LNs and peak hyperpolarization for Cluster 1 LNs (orange). *On the right,* a histogram depicting the timing of initial depolarization peaks for both clusters.

**Supplementary Figure 2.**
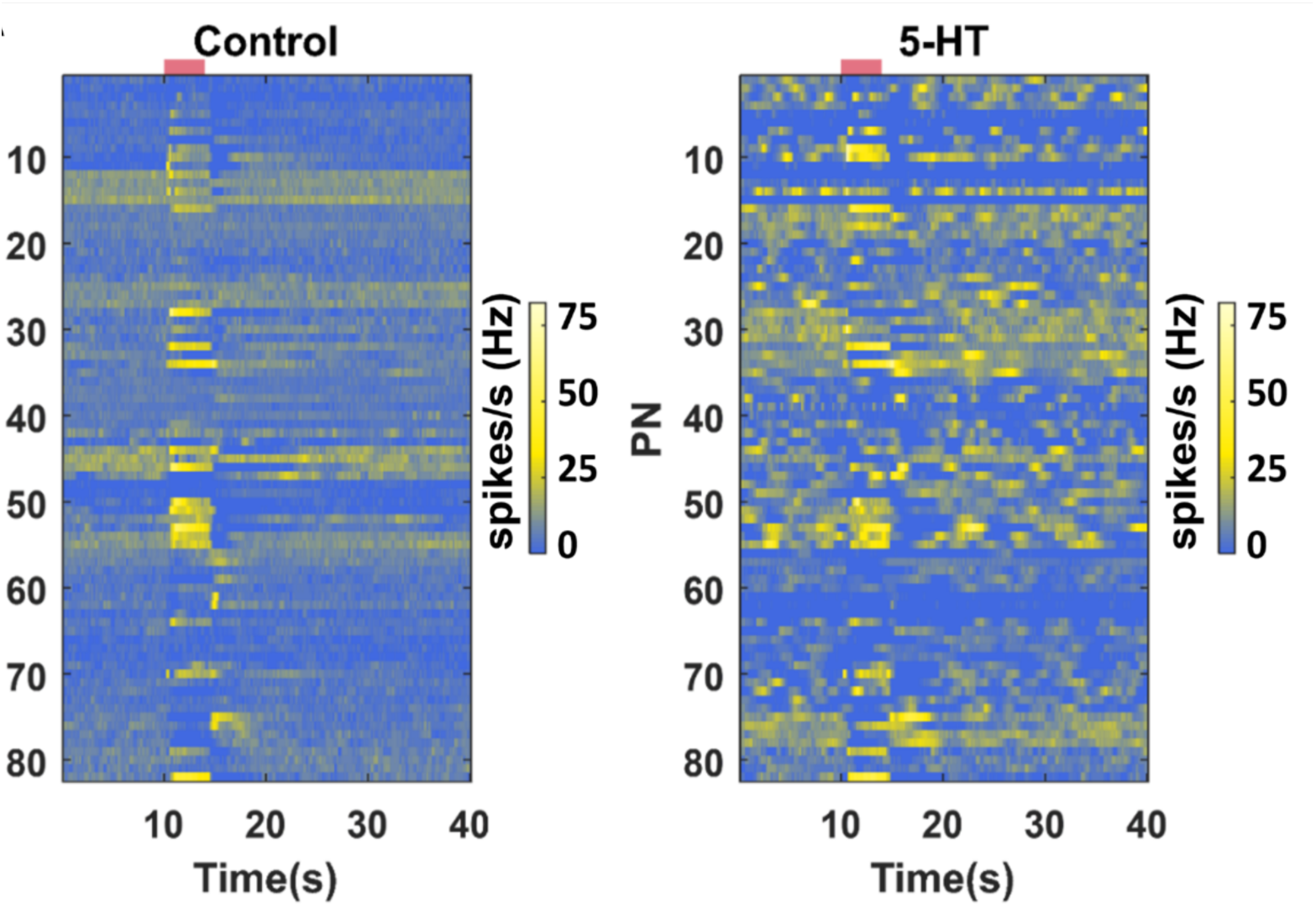
Serotonin changes spontaneous activity of PNs to bursting. Trial-averaged spiking activity as a function of time is shown for 82 neurons. The hotter color identifies the higher average firing rates per bin (200 ms). Each row represents one PN, each column represents one-time bin. The red bar identifies the odor presentation time period. The heatmap on the left shows the PN activity matrix before 5HT application and the heatmap on the right shows the PN activity matrix (neurons sorted in the same order) after 5HT exposure*. Data replotted from Bessonova et al*.

**Supplementary Figure 3.**
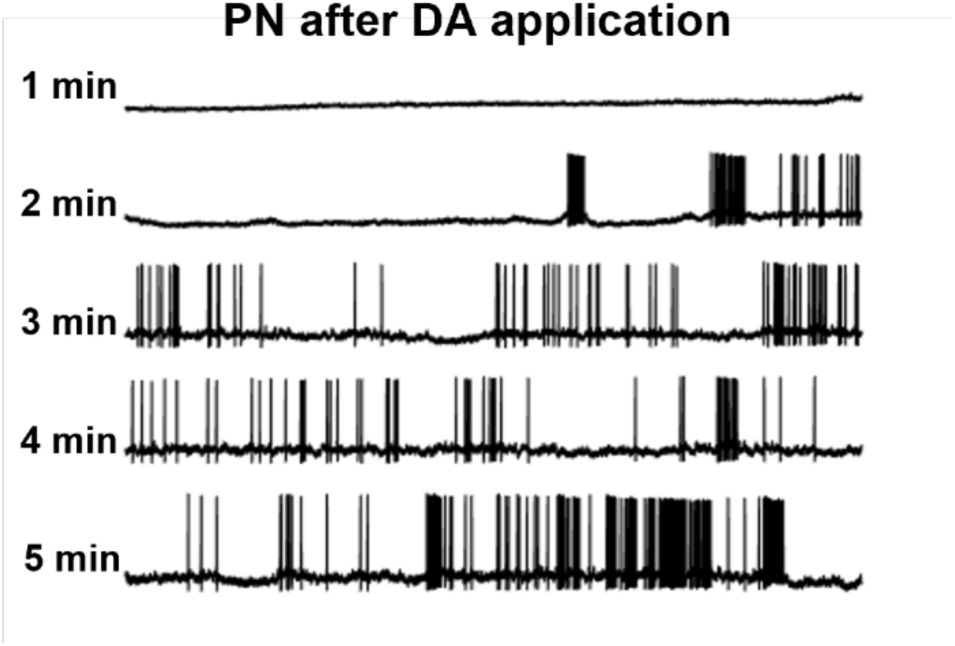
Dopamine completely shuts down PN activity immediately after application. A representative intracellular recording displays the membrane potential over time for a single projection neuron in the locust antennal lobe after application of dopamine. Each row represents a recording after one minute.

**Supplementary Figure 4.**
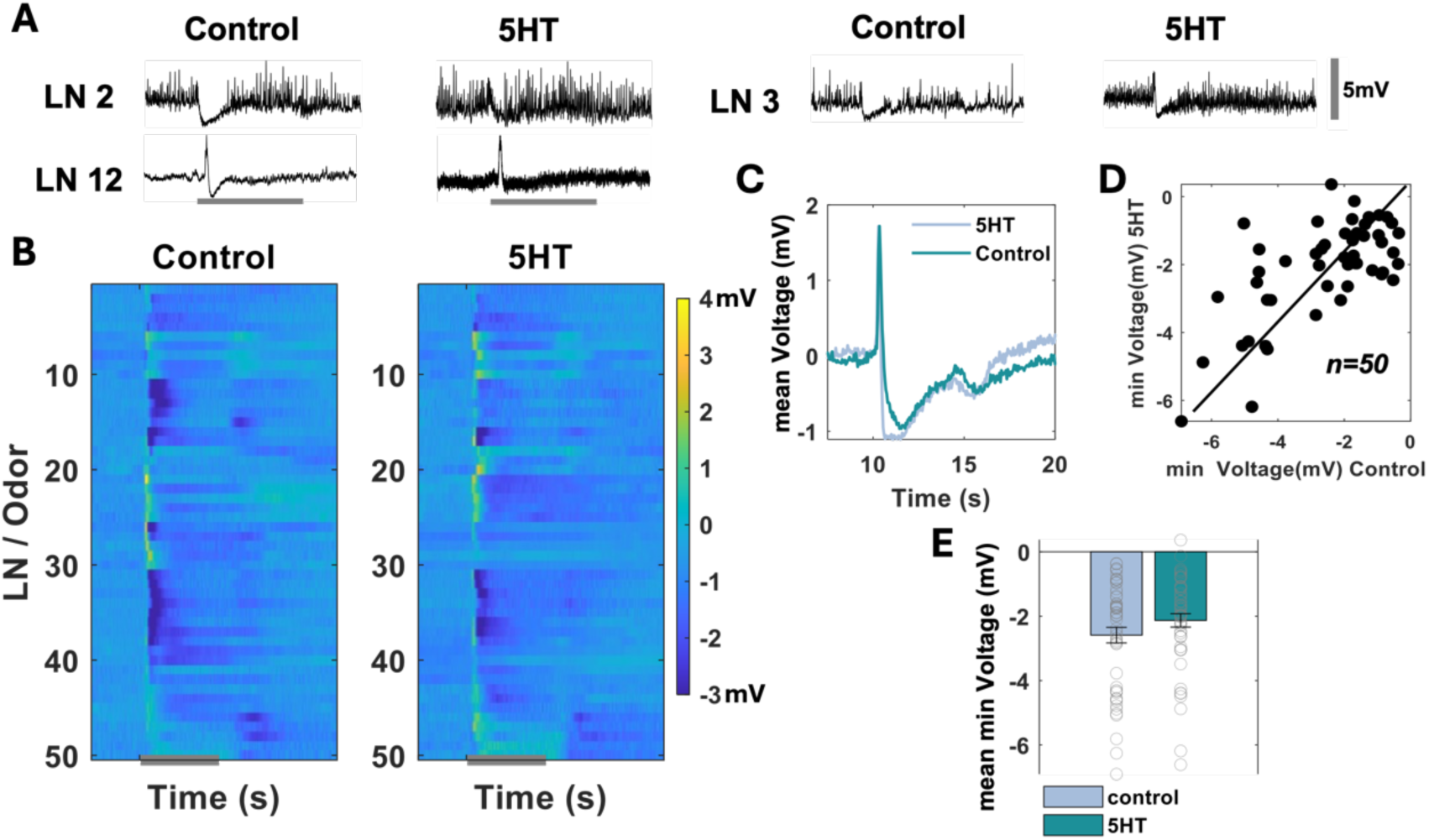
Local neuron activity is not altered after exogenous serotonin introduction. **(A)** Representative intercellular recording shows membrane potential as a function of time for three distinct local neurons (LNs) before (control) and after application of serotonin. No significant change in odor-evoked response was observed. The vertical grey scale bar represents 5 mV, and the horizontal grey bar identifies an odor presentation period. **(B)** Trial-averaged voltage values as a function of time are shown for LNs before (control) and after treatment with serotonin. Responses to five odorants are shown (hexanol, benzaldehyde, linalool, citral, and geraniol). The raw membrane potential in 200-ms time windows was averaged and baseline-subtracted before being shown here as a heat map. Each row corresponds to one trial, and the average of five trials represents one LN- odor combination. The hotter color identifies higher value (depolarization), and the lighter color indicates lower value (hyperpolarization). The horizontal grey bar indicates the odor presentation period. The scale bar represents the baseline-subtracted, binned averaged voltage in millivolts *(mV)*. The grey bar identifies the 4 seconds of odor presentation. **(C)** PSTHs of all LN/odor combinations are shown before (lighter shade color) and after neuromodulator (darker shade color) application. **(D)** Minimum voltage value during odor presentation in the control condition and after neuromodulator application for each LN/odor combination. **(E)** The mean minimum voltage value across LNs during odor presentation in the control condition (lighter bar) and after neuromodulator application (darker bar) is compared. Grey circles showcase a single minimum value for all LN/odor combinations. Error bars indicate SEM. Significant differences, if present, are identified in the plot (standard paired sample t-test, *p<0.05; **p<0.01; ***p<0.001).

**Supplementary figure 5.**
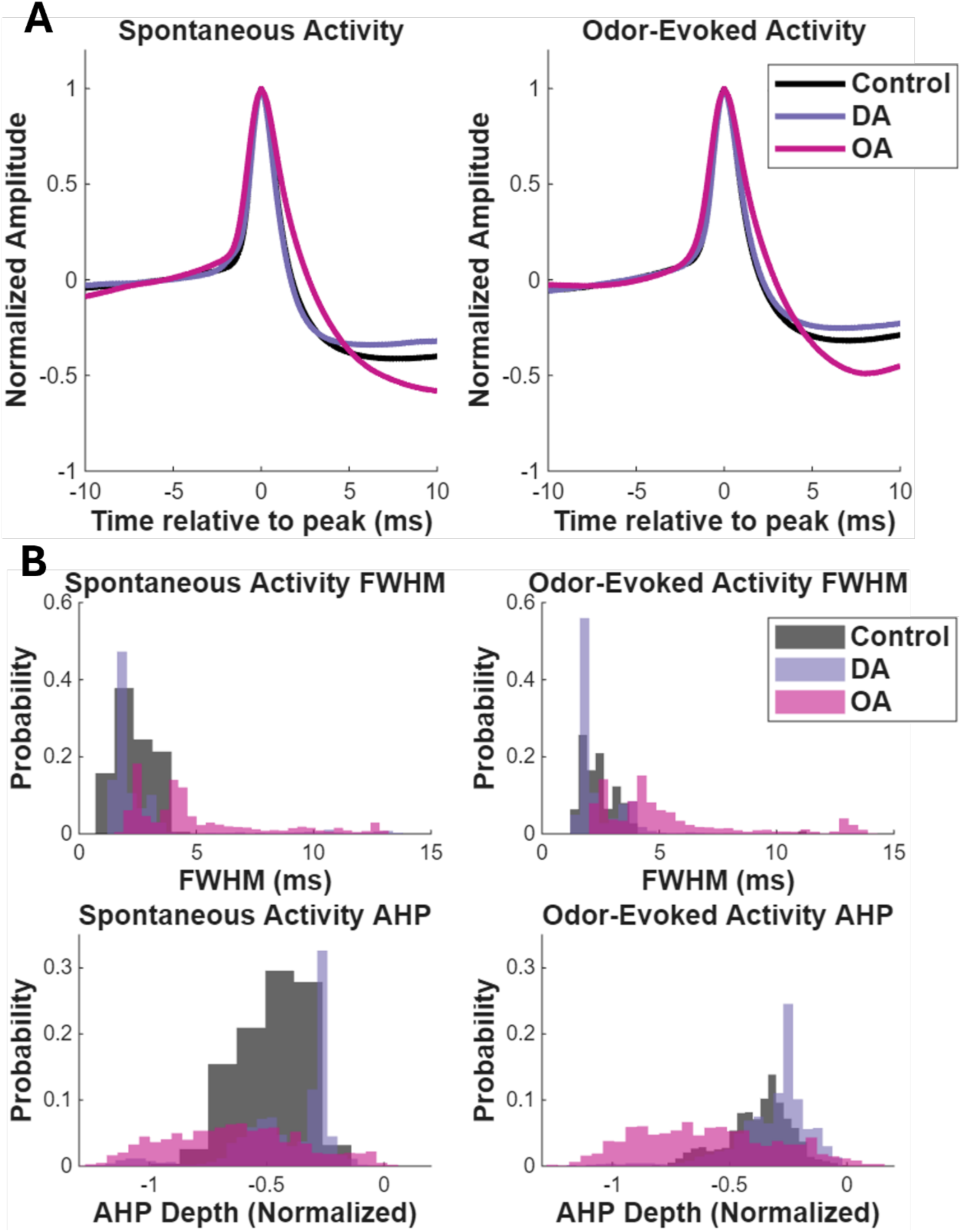
Octopamine alters action potential waveform shape while dopamine produces minimal effects. **(A)** Mean action potential waveforms (±SEM) for PNs before (control, black) and after dopamine (purple) or octopamine (pink) application during spontaneous (left) and odor-evoked (right) activity. Time is shown relative to the action potential peak (0 ms). All action potentials were normalized to unit amplitude based on peak amplitude. (Control: n=17,952 spikes from 20 PNs; Dopamine: n=4,811 spikes from 12 PNs; Octopamine: n=2,349 spikes from 8 PNs) **(B)** Histogram distributions of action potential full-width at half-maximum (FWHM, top) and afterhyperpolarization depth (AHP, bottom) for spontaneous (left) and odor-evoked (right) activity. Histograms show normalized probability distributions. AHP depth measured as the minimum voltage within 30 ms following the action potential unit peak.

## Notes

https://github.com/b0r2023/Distinct-mechanisms-mediate-dopamine-octopamine-opponency-in-an-insect-model-of-olfaction

### Competing Interest Statement

The authors have declared no competing interest.

